# Comparing complex variants in family trios

**DOI:** 10.1101/253492

**Authors:** Berke Ç. Toptaş, Goran Rakocevic, Péter Kómár, Deniz Kural

## Abstract

**Motivation:** Several tools exist to count Mendelian violations in family trios by comparing variants at the same genomic positions. This naive variant comparison however, fails to assess regions where multiple variants need to be examined together, resulting in reduced accuracy of existing Mendelian violation checking tools.

**Results:** We introduce VBT, a trio concordance analysis tool, that identifies Mendelian violations by approximately solving the 3-way variant matching problem to resolve variant representation differences in family trios. We show that VBT outperforms previous trio comparison methods by accuracy.

**Availability:** VBT is implemented in C++ and source code is available under GNU GPLv3 license at the following URL: https://github.com/sbg/VBT-TrioAnalysis.git

**Supplementary information:** Supplementary materials are available at *Biorxiv*.

## 1. Introduction

Recent technological advancements enabled a rapid progress in our understanding and characterization of the human genome, assessment of the scale and extent of genomic variation present in human genome (The 1000 Genome Project Consortium et al., 2015) as well as creation of a vast body of knowledge related to the functioning of human body and rare diseases (Jamuar et al., 2015). Pedigree-based genetics plays a crucial role in uncovering the genetic origins of diseases, where family trios are analyzed and genomic variants that disagree with Mendel’s law of segregation are identified.

A particularly important use case of trio analysis is the identification of de novo mutations which have repeatedly been implicated in rare and complex diseases (Hidalgo *et al.*, 2016; Deciphering Developmental Disorders Study, 2017). De novo mutations occur with relatively low frequencies (1.2x10^-8^) (Conrad *et al.*, 2011; Kong *et al.*, 2012) compared with average amount of variants a person has. Therefore, accurate strategies are essential for identification of such variants which typically starts with assessing Mendelian Inheritance rules of the calls from family trios followed by using sophisticated statistical models (DeNovoGear (Ramu *et al.*, 2013), PhaseByTransmission (Francioli *et al.*, 2016)).

Such trio analysis can also be used for truth-free benchmarking of variant calling pipelines (Douglas *et al.*, 2002; Pilipenko *et al.*, 2014; Nutsua *et al.*, 2015; Kómár *et al.*, 2017). Due to the very small mutation rate in human genome, all Mendelian violations can be considered as sequencing/variant calling errors. Trio concordance analysis is useful where no truth-set exists and allows using variants from all regions of the genome as opposed to current whole genome truth-sets which are limited to a set of high confidence regions in a few samples, excluding many regions of the genome (Zook *et al.*, 2014). Improved truth-free benchmarking will also guide the development of future genome analysis pipelines such as graph genome pipeline (Rakocevic et al., 2017).

Several tools exist (RTG-mendel, GATK-SelectVariants, Vcftools-mendel) that count Mendelian violations using naive loci-by-loci variant comparison. In this approach, each record in the merged trio vcf is processed independently, and only variants with coinciding reference positions are analyzed together. This method fails to provide an accurate analysis in cases where multiple records are affecting the same locus.

Here we address a problem during the identification of Mendelian violations in the data from a family trio, one which arises from varying variant representations. Regions with several overlapping variants often have a number of different ways in which they can be represented, all of which conform to the widely accepted VCF standard (Danecek *et al.*, 2011); the same is true for most variants which are complex in nature, and even some simple indels (Figure 1a). The choice of which of the possible representations is produced often depends on the variant context (other nearby variants) and the set of sequencing reads used to identify the variant. If this choice happens to be different between different members of the pedigree, comparing the three sets of calls position by position will result in detection of Mendelian violations, even though the underlying haplotypes are Mendelian compliant (Figure 1b).

**Figure 1:**
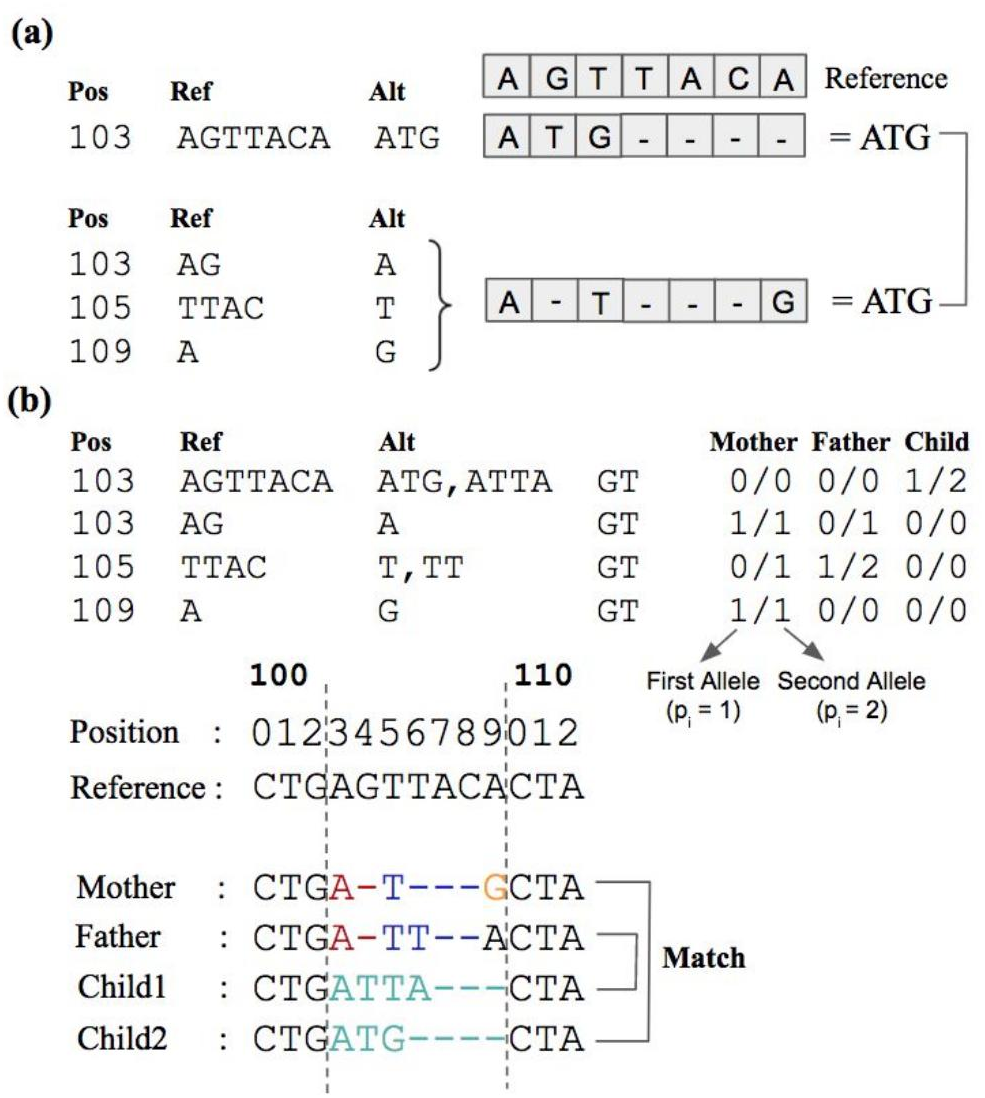
**(a)** Representation difference in indels. Variant in position 103 is represented as a single indel in first vcf and 2 indels + 1 SNP in second vcf. After they are applied on the reference sequence, it is seen that they are equivalent. **(b)** A toy example of variant representation difference in family trios. Naive trio comparison tools marks all 4 records as Mendelian violation. However a consistent combination can be found if they are processed together.

Problems related to variant representation has been recognized in the context of benchmarking NGS data processing methods, and numerous approaches have been developed for comparing two sets of results for a single sample (SMaSH(Talwalker *et al.*, 2014), Vcfeval(Cleary *et al.*, 2015), VarMatch(Sun *et al.*, 2017)). However, none of these tools are capable of resolving the issue with data from a family trio.

One way to unify variant representations across a family trio is to use a joint variant caller such as GATK GenotypeGVCFs. This tool aggregates variants of multiple samples by combining overlapping trio records and re-genotyping them. Although joint calling resolves many representation issues, it is still unable to merge complex overlapping indels affecting the same site. This method also requires GATK (McKenna *et al.*, 2010) HaplotypeCaller as variant caller which eliminates the benchmarking purpose of trio analysis.

In this paper, we present VBT, a Mendelian violation detection tool that uses advanced variant comparison to deal with ambiguities arising from different variant representations. VBT extends the variant comparison algorithm of vcfeval (Cleary *et al.*, 2015) for trio concordance analysis. We show that VBT outperforms all previous trio comparison methods in terms of accuracy of detecting Mendelian violations.

## 2. Methods

### 2.1 Variant Comparison and Ideal Mendelian Equation

For a diploid variant set V from a single sample VCF, we define the phasing vector, 
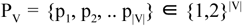
, where the i-th value (1 or 2) indicates whether the first or second allele (Figure 1b) of the i-th variant is selected for the maternal haplotype. Similarly P_v’_ denotes the opposite phasing vector 
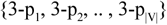
 which indicates the alleles on the paternal haplotype, not selected by P_v_. A haplotype function h(V, P_V_) is defined (Cleary *et al.*, 2015) to produce the haplotype sequence obtained by applying all variants of V to the reference sequence using the P_v_ phasing vector. vcfeval defines the variant matching problem as finding the optimal sets of variants X^opt^
, Y^opt^, and their corresponding phasing vectors 
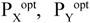
, that solves the following optimization problem:

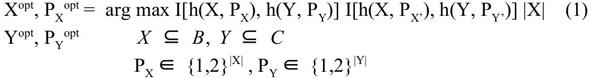

where B and C denote baseline (gold standard) and called (test) variant sets, and X^opt^ and Y^opt^ are the sets of variants which maximize the number of matching variants in baseline and called variant set. I[seq1, seq2] is the indicator function that is 1 if seq1 = seq2, and 0 otherwise.

For VBT, we aim to extend the definition in eq. (1) for family trios to detect Mendelian violations. Let M, F and C represent the sets of variants of mother, father and child, respectively. We define the trio matching problem as finding the optimal sets X^opt^, Y^opt^, Z^opt^ and their corresponding phasing vectors 
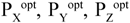
 that solve the following optimization problem:

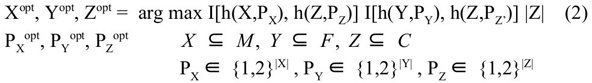

Eq. (2) maximizes the number of child variants that follow Mendelian Inheritance rules. Z^opt^ denotes the set of Mendelian-consistent variants in the child, and the remaining child variants C \ Z^opt^ are marked as Mendelian violations.

During variant comparison in VBT and vcfeval, *syncpoints* (Cleary *et al.*, 2015) are calculated procedurally which are genomic positions that delimit regions where variants within the region can be processed independently. For the variants inside a single region (subset), there are 3^|subset|^ combinations where, for each family member and each variant r, there are three possibilities: excluding r, including r with p_r_ = 1, or including r with p_r_= 2. Therefore the time and space complexity of the fundamental algorithm of vcfeval scales exponentially with the size of the typical subset. |subset| is mostly small (ie. < 5) when comparing two VCF files. For larger subsets, even though some pruning strategies are applied to keep the total variant combination low, vcfeval uses a cutoff strategy by skipping regions where the total combination count exceeds the predefined limit.

In trio comparison with eq. (2), subset sizes are increased since we have mother, father and child variants in subsets instead of variants from two VCFs. In addition, for trio comparison, region intervals tends to be larger due to less number of *syncpoints*. For these two reasons, a similar cutoff strategy (estimated as |subset| > 15 with pruning strategies) using eq. (2) would cause to skip more than 200,000 variants (out of ~18 million single sample variants combined) as seen in Figure 2. Instead of using eq. (2) directly in VBT, we use a heuristic, described below, that alleviates exponential scaling at the cost of a small chance of making mistakes.

**Figure 2:**
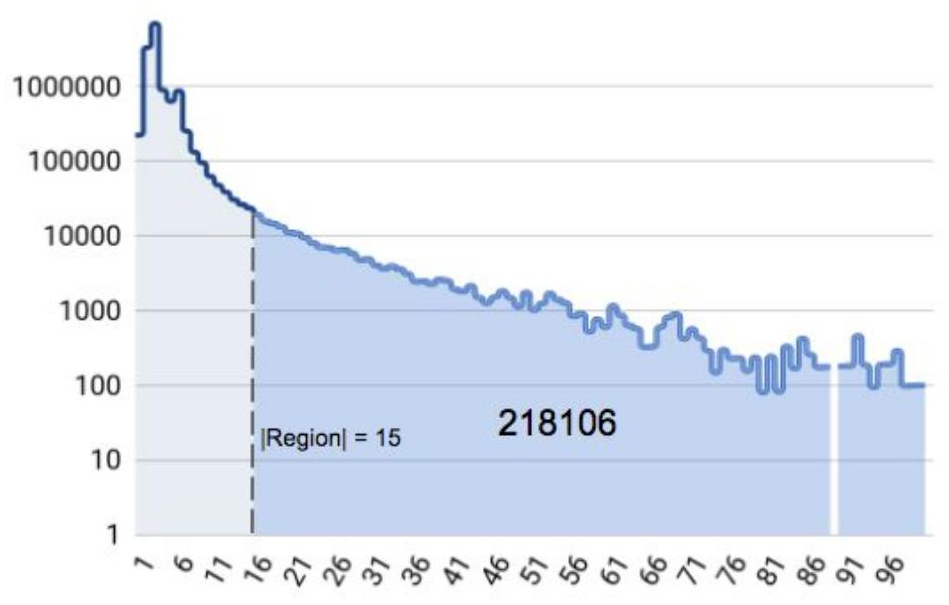
Number of total non-ref variants vs region size for Central European (CEU) Trio using HaplotypeCaller. Regions are obtained using intermediate results (syncpoints) of vcfeval in ‐‐squash-ploidy mode. Mother-child and father-child VCFs are compared separately and resulting two sync point sets are intersected.

### 2.2 Search Space Reduction and Same Allele Match Elimination

We aim to decrease the number of possible combinations by separating the two indicator functions in eq. (2) for mother-child and father-child variant sets, and optimize the maternal and paternal haplotype sequences of child separately using the following equation:

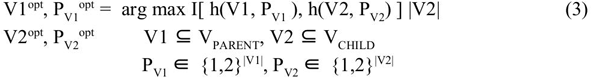

Where V2^opt^ is the set of child variants that shares an allele with parent V_PARENT_. After separate processing, we can mark the child variants that exists in both haplotype sequences, and the remaining child variants become Mendelian violations. Note that Eq. (3) is not the same as the duo comparison method of Eq. (1). The former requires matching both genotypes, while the latter requires matching one only.

When processing mother-child and father-child variants separately, we need to guarantee that the child’s haplotypes use opposite phases P_Z_ and P_Z’_. For heterozygous child variants, if one of the two alleles is not present in either the mother-child or the father-child sequences, they should be reported as a Mendelian violation. For instance, if genotype of mother is A/A, father is C/A and child is A/G for a multi-allelic SNP variant at the same position, then the child’s variant matches with both parents’ variants with allele A. Allele G of child on the other hand is not present in any of the parents. Although pairwise comparisons with both parents indicate one matching allele of the child, this locus is a Mendelian violation because the same phase is used for both matches. Inmost cases, we can indeed mark it as such. We call this condition *same allele matching*.

In a family trio, child variants often match to parent variants with both of their alleles. For these child variants, any of the two alleles can be present in the final haplotype sequence, and at this stage we select the phase arbitrarily. If, after pairwise comparison, the same allele of the child is matched with both parents, we would mark it as violation, following the rules of same allele matching condition, even though, in this case, we could flip the phasing with one parent without breaking the match and resolve the inconsistency. To identify these variants, we apply the duo comparison function (ie. Eq. (1)) to mother-child and father-child variants:

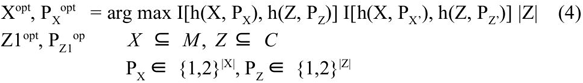

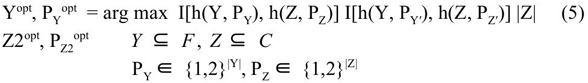

where Z1 and Z2 are child variants sharing both alleles with mother and father variants respectively. From the remaining variants M \ X^opt^ ( =: MM), F \ Y^opt^ ( =: FF), C \ Z1^opt^ ( =: CC1) and C \ Z2^opt^ ( =: CC2); we obtain all child variants sharing a single allele by maximizing the number of variants in constructing a single haplotype sequence, ignoring the alternative phases of variant sets:

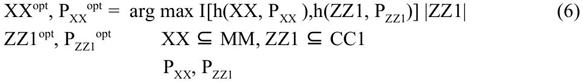

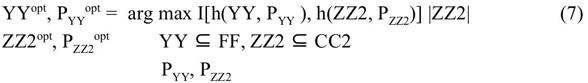

where, during maximization, P_XX_, P_YY_, P_ZZ1_ and P_ZZ2_ are required to be such that the reference allele (‘0’) is never used in any comparison. I.e. if a variant has the genotype 1|0, the corresponding phasing is not allowed to take the value 2, because that would correspond to the ‘0’ allele.

The reason for not allowing the reference allele to be used is the ambiguity caused by identical representation of excluded variant and included reference allele. If we allow reference alleles in the haplotype function for eq. (6) and (7), child variants having reference phasing would always be included regardless of the corresponding parent variant. For example, if genotype of mother is 0/2, father is 2/2 and child is 0/1 for a variant, mother and child variants would be included in mother-child side because they share ‘0’ genotype. In father-child side, there is no shared allele but once the father variant is excluded, that position becomes reference and child variant alone could be included with ‘0’ genotype. At the end, child variant would be present on both mother and father final haplotypes and would be marked as Mendelian consistent, while it is a violation in reality. With the above restriction on the phasing vectors, we eliminate this mistake.

Using eq. (4), (5), (6) and (7), we construct the VBT pipeline as shown in Figure 3 to obtain our four child variant set Z1^opt^, Z2^opt^, ZZ1^opt^, ZZ2^opt^ with their phasing information 
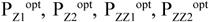
. Using these 4 variant set, we check how many of them exist in both mother-child and father-child side to determine Mendelian violations with Algorithm 1:

**Figure 3:**
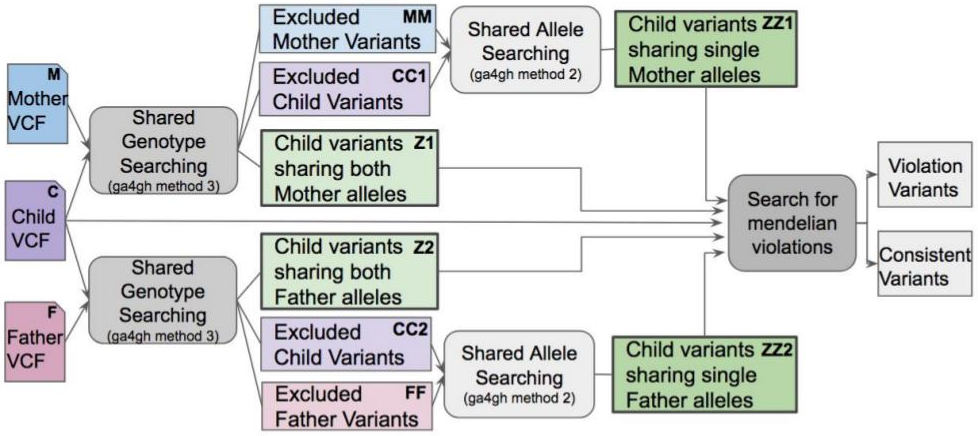
VBT pipeline using vcfeval best path algorithm and GA4GH benchmarking standard methods(Zook, 2015). Included variants are present in the best common path between parent and child while excluded variants are eliminated from that path.

**Algorithm 1:**
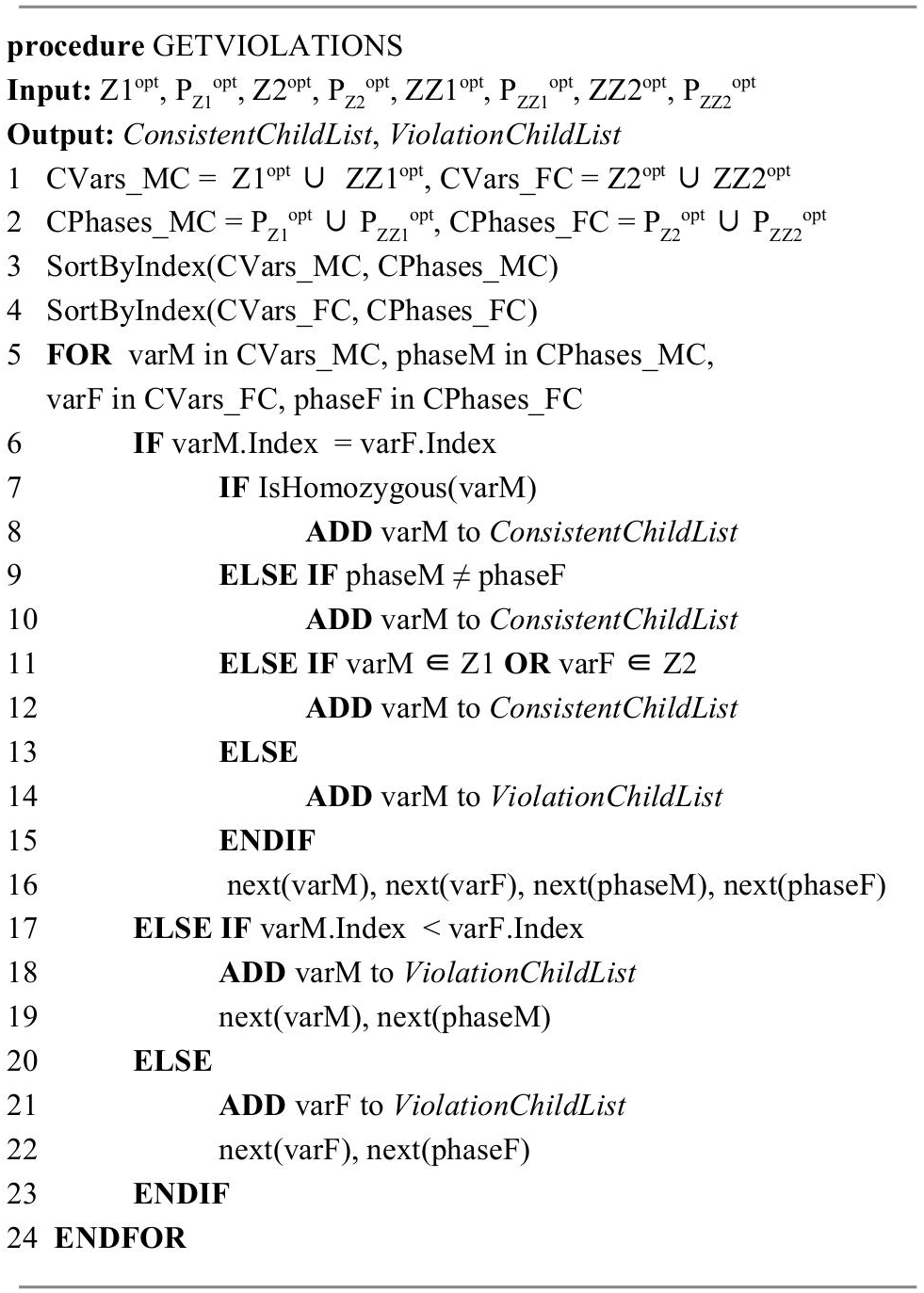
Same Allele Match Elimination

In Algorithm 1, in order to identify child variants that share an allele with both parents, we first merge the shared genotype and shared allele child variant sets keeping the information of belonging sets for each variant at lines (1) and (2) of the pseudocode. Then we sort the merged child variant sets by variant indexes (order in VCF) at lines (3) and (4). At line (7), we check the condition where child variant is homozygous and same allele matching condition is ignored. At line (9) we check whether heterozygous child variants match with parents with different phasings. At line (11) we check if child variant matches to parent with both alleles so that alternative phasing can also be used to avoid same allele matching condition. We use *next* command to iterate to the following variant/phase at line (16), (19) and (22). At the end, we obtain the list of Mendelian violations and consistent child variants for the input sets Z1^opt^, Z2^opt^, ZZ1^opt^ and ZZ2^opt^.

In eq. (6) and (7), reference alleles of child variants are ignored during maximization calculation. As a result, child variants that are matching one parent with non-reference allele and other parent with reference allele are marked as Mendelian violation after processing variants with Algorithm 1. In order to identify and correct the decision of these child variants, we use the following equations:

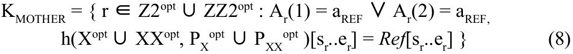

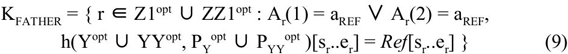

Where s_r_ and e_r_ denote the start and end position of variant r and *Ref* denotes the reference sequence string. Indexing of the haplotype sequence is inherited from the reference sequence. A_r_(k) is the allele function that represents the allele of variant r with the phase selection of k ∈ {1,2} and a_REF_ is the reference allele of variant. K_MOTHER_ and K_FATHER_ are the sets of consistent child variants that shares a reference allele with Mother and Father variants respectively and they are inserted to the set of consistent variants. The remaining unprocessed child variants (ie. C \ (Z1^opt^ ⋃ Z2^opt^ ⋃ ZZ1^opt^ ⋃ ZZ2^opt^)) are inserted to the set of violations.

Once we obtain decisions of all child variants, we merge mother, father and child VCF as a trio by merging variants at the same position. Then, we apply three post-processing steps on the merged VCF:

1. Assign Mendelian decision to sites where child has no variant (ie. homozygous ref child variants in merged trio). For each hom-ref child variant, final haplotype sequences of both mother and father are checked. If none of the parent haplotype sequences is equal to reference at the child variant’s location, then the variant is marked as violation.
2. Consolidate the decision for variants affecting the same position in the final haplotype sequence. Consistent VCF record decisions are changed to violation if there is at least one overlapping violation VCF record.
3. Exclude sites where *nocall* is reported by at least one family member. *Nocall* variants are sites where insufficient information is available to determine genotypes and they are usually represented as ‘./.’ at genotype column of VCF records.

Finally, we solve the maximization problem given in eq (4), (5), (6) and (7) using dynamic programming solution of Cleary *et al.*, (2015) by slightly changing the algorithm for reference overlapping variants which is described in Supplementary text, Section 2.

### 2.3 Evaluation Methods and Data

A truth set for trio analysis does not exist for direct result comparison. Instead, we use alternative testing methods to compare VBT and existing tools. For our experiments, we use high coverage alignments of Central European (CEU) individuals (NA12878, NA12891 and NA12892) available at 1000 genomes phase 3 ftp server (ftp://ftp.1000genomes.ebi.ac.uk/vol1/ftp/phase3/data/).

For our first testing experiment, we construct trios from single individual samples by changing their variant representations. We use FreeBayes (Garrison *et al.*, 2012) to generate unnormalized VCF files. Then we use Vt norm (Tan *et al.*, 2015) to alter variant representations of VCF. By using vcftools merge, we merge two identical unnormalized VCFs (playing the roles of mother and father samples) and one normalized VCF (playing the role of child sample). Since all trio samples belongs to the same individual, we expect to see zero Mendelian violations by all Mendelian violation checking tools.

For the second testing experiment, we implement a Mendelian violation validator that checks all possible combinations of variant phasings in a given set of small regions. We obtain the regions by merging *syncpoints* yielded by mother-child and father-child variant comparisons. Our validation pipeline performs a comparative analysis of unique Mendelian violations between two Mendelian violation checking tools. That is why, we first select the regions where either VBT or naive tools find a Mendelian Violation. We ignore the regions in which both tools found no Mendelian violations.

For each selected region, we discard the variants that are marked as Mendelian violation and, using the remaining variants, we seek for a Mendelian consistent combination. If no consistent combination can be found, the region is marked as *missing MV* for that tool. If a consistent combination can be found for both VBT and the naive method for a region, then we compare the reported violation counts for that region in order to check whether there are extra Mendelian violations reported. We accept the decision of tool with less Mendelian violation as correct and mark that region for the other tool as *extra MV*. A more detailed overview of our validation pipeline can be found in Supplementary Text, Section 3.

## 3. Results

VBT resolves variant representation differences in family trios efficiently by maximizing matching child variants with mother and father separately instead of using the ideal trio comparison function (eq 2). This enables covering nearly all regions in datasets and provide VBT a reasonable running time, which varies between 6 and 8 minutes on Amazon c4.4xlarge instance (Intel Xeon E5 2.9 Ghz, 16 vCPU, 30 GiB Memory) depending on complex region count for whole human genome that contains 6.1 million vcf record.

In our first test scenario, we use the trios we generated from a single CEU sample to show that naive trio comparison tools produce wrong Mendelian decisions due to variant representations. We compared VBT, naive (line-by-line) Mendelian error checking tools (RTG-mendel, GATK-SelectVariants, Vcftools-mendel) and PhaseByTransmission (PBT). For PBT, we used both 2x10^-2^ and 10^-8^ as mutation rates, and obtained the same number of *corrections* plus *mutations*. As seen in Table 1, VBT correctly outputs no violations for all three test data while the other tools output more than seventy thousand violations.

**Table 1.** Violation counts of different tools where the input trio is constructed from a single sample with different variant representations

| Input Sample | VBT | Naive | PBT |
| --- | --- | --- | --- |
| NA12878 as trio | 0 | 76,867 | 72,111 |
| NA12891 as trio | 0 | 78,854 | 73,396 |
| NA12892 as trio | 0 | 76,176 | 72,674 |

Naive check tools include RTG -mendel, GATK -SelectVariants and Vcftools -mendel. For PhaseByTransmission(PBT), sum of violation count and corrected genotype count is used in this table.

In the second experiment, we used CEU samples to compare trio concordance rate of different variant callers, FreeBayes (fb), UnifiedGenotyper (ug) and HaplotypeCaller (hc). In addition, we apply normalization (vt norm) on FreeBayes outputs and add it as a fourth testing set to see if normalization can reduce errors of naive comparison tools. As a final testing trio, we produce gVCFs using HaplotypeCaller and jointly call them using GATK GenotypeGVCFs. We used vcftools (v0.1.14) to merge VCF files of individual samples except the jointly called HaplotypeCaller trio VCF.

After we generate 5 trio VCFs using different variant calling pipelines, we ran VBT and naive checking tool for each trio and compare their result using our Mendelian violation pipeline. Table 2 shows the numbers of total violations, falsely identified violations and missed violations for the two methods, and for the 5 different variant calling pipelines. For all 5 testing pipeline, VBT has over 99% precision and recall values (Figure 4). The precision of naive tools and VBT is closer for HaplotypeCaller and jointly-called HaplotypeCaller because the representations of called variants are more similar across the samples compared to other variant callers.

**Table 2.** Violation validation results of different variant calling pipelines using CEU trio bam files aligned with BWA-MEM.

|  | VBT |  |  | Naive |  |
| --- | --- | --- | --- | --- | --- |
|  | Total MV Regions | Extra MV Regions | Missing MV Regions | Extra MV Regions | Missing MV Regions |
| **fb** | 163,094 | 446 | 876 | 21,313 | 18,656 |
| **ug** | 120,709 | 383 | 905 | 6,475 | 17,475 |
| **hc** | 73,501 | 634 | 353 | 32 | 6,560 |
| **fb+norm** | 163,606 | 407 | 1,244 | 23,341 | 15,859 |
| **hc joint** | 39,488 | 334 | 84 | 226 | 1,643 |

Comparison results of FreeBayes (fb), HaplotypeCaller (hc) and UnifiedGenotyper (ug). Autosomes only. No filtration is applied to the data. Regions may contain zero or multiple Mendelian violations.

**Figure 4:**
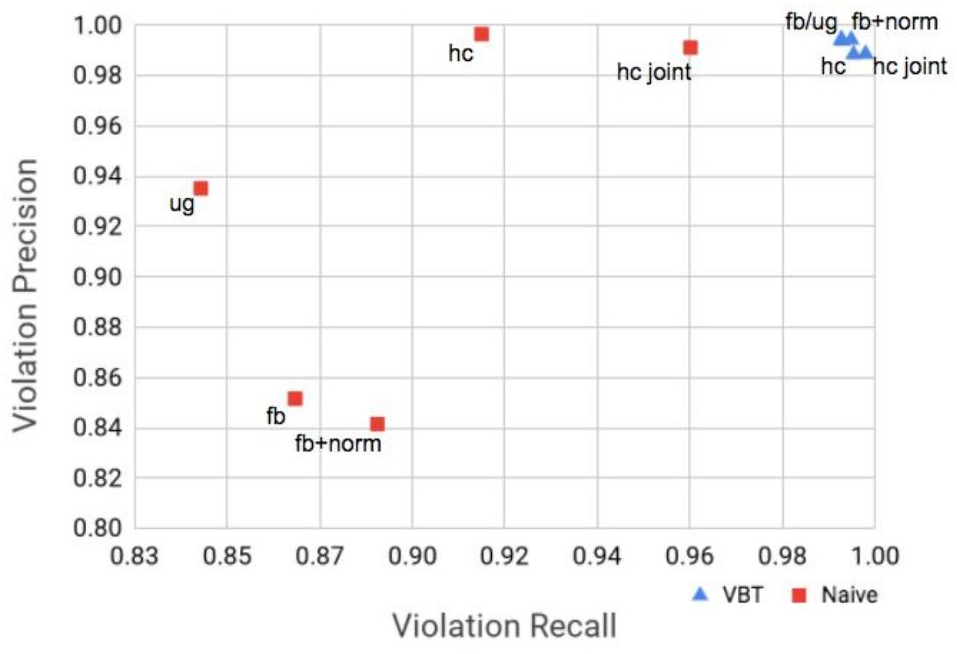
Violation Precision vs Recall plot of VBT and Naive tools for 5 different variant calling pipelines. Recall is defined as *(Total MV Region - Extra MV - Missing MV) / (Total MVRegion - Extra MV)* and precision is defined as *(Total MV Region - Extra MV - Missing MV) / (Total MV Region - Missing MV)* according to Table 2.

## 4. Discussion

In this work, we presented VBT, a mendelian violation detection tool that is capable of comparing complex indels in family trios. We showed with our test scenarios that, VBT has better accuracy than the existing tools.

With our method, we process mother-child and father-child haplotype chains separately. However obtained local best paths from mother-child and father-child duos are not always identical to the global optimum of the trio. This introduces a small error rate visible on Table 2. With our proposed method, we accept a small loss in accuracy in exchange for being able to analyze whole genome data with minimum number of skipped regions plus we gain a significant performance by reducing the search space.

Instead of our method, ideal Mendelian function (eq. 2) could be implemented to minimize error rate by skipping variants in more complex regions (~1% of all records). Then, skipped variants could either be ignored, be processed using naive variant comparison or be processed using the current VBT strategy.

VBT’s accuracy can further be improved by correcting wrong/missed decisions by comparing vcf output with naive comparison as a post-processing step. In regions where the naive method and VBT disagrees, a nonlinear violation check can be performed by generating all possible subsequences for that region, similarly to our violation validation pipeline. This would not increase overall running time considerably because the slow nonlinear checking method would be invoked only for regions where the naive method disagrees with VBT. As a result, VBT would serve as a cost-efficient detector informing us whenever the naive comparison methods are not enough.

VBT does not alter variant representations during or after variant comparison. Instead, we keep the original variant representations and add additional info tag that whether a variant is Mendelian violation or not. The advantage of this is the ability of tracking variants for benchmarking purposes. The disadvantage is that existing tools which require trio analysis such as PhaseByTransmission are not able to use VBT output directly and need to read Mendelian decision annotation from output vcf records.

## 5. Acknowledgements

We would like to thank Maxime Huvet, Maria C. Suciu and Amit Jain for trio benchmarking discussions and thank Sun-Gou Ji, Yilong Li, Morten Källberg, Gülfem Demir, James Spencer and Kaushik Ghose for their valuable comments on VBT.

